# Inherited *MUTYH* mutations cause elevated somatic mutation rates and distinctive mutational signatures in normal human cells

**DOI:** 10.1101/2021.10.20.465093

**Authors:** Philip S. Robinson, Laura E. Thomas, Federico Abascal, Hyunchul Jung, Luke M.R. Harvey, Hannah D. West, Sigurgeir Olafsson, Bernard C. H. Lee, Tim H.H. Coorens, Henry Lee-Six, Laura Butlin, Nicola Lander, Mathijs A. Sanders, Stefanie V. Lensing, Simon J.A. Buczacki, Rogier ten Hoopen, Nicholas Coleman, Roxanne Brunton-Sim, Simon Rushbrook, Kourosh Saeb-Parsy, Fiona Lalloo, Peter J. Campbell, Iñigo Martincorena, Julian R. Sampson, Michael R. Stratton

## Abstract

Cellular DNA damage caused by reactive oxygen species is repaired by the base excision repair (BER) pathway which includes the DNA glycosylase MUTYH. Inherited biallelic *MUTYH* mutations cause predisposition to colorectal adenomas and carcinoma. However, the mechanistic progression from germline *MUTYH* mutations to **M**UTYH-**A**ssociated **P**olyposis (MAP) is incompletely understood. Here, we sequenced normal tissue DNAs from 10 individuals with MAP. Somatic base substitution mutation rates in intestinal epithelial cells were elevated 2 to 5-fold in all individuals, except for one showing a 33-fold increase, and were also increased in other tissues. The increased mutation burdens were of multiple mutational signatures characterised by C>A changes. Different mutation rates and signatures between individuals were likely due to different *MUTYH* mutations or additional inherited mutations in other BER pathway genes. The elevated base substitution rate in normal cells likely accounts for the predisposition to neoplasia in MAP. Despite ubiquitously elevated mutation rates, individuals with MAP do not display overt evidence of premature ageing. Thus, accumulation of somatic mutations may not be sufficient to cause the global organismal functional decline of ageing.

**Summary**

## INTRODUCTION

The genomes of all normal human cells are thought to acquire mutations during the course of life. However, the mutation rates of normal cells and the processes of DNA damage, repair and replication that underlie them are incompletely understood^1–8^. A ubiquitous source of potential mutations is DNA damage caused by reactive oxygen species (ROS) which are formed as by-products of aerobic metabolism^9^. ROS cause a variety of DNA lesions, the most common being 8-oxoguanine (8-OG)^10^. As a consequence of mispairing with adenine during DNA replication, 8-OG can cause G:C>T:A (referred to as C>A for brevity) transversion mutations^11^. Under normal circumstances, 8-OG and its consequences are efficiently mitigated by the Base Excision Repair (BER) pathway effected by DNA glycosylases; oxoguanine DNA glycosylase (OGG1) removes 8-OG^12^ and MutY DNA glycosylase (MUTYH) removes adenines misincorporated opposite 8-OG^13^.

Mutations in *MUTYH* engineered in experimental systems can impair its glycosylase activity, reducing its ability to excise mispaired bases and leading to an increased rate of predominantly C>A mutations^14–18^. *MUTYH* mutations inherited in the germline in humans cause an autosomal recessive syndrome (MUTYH-associated polyposis, MAP) characterised by intestinal adenomatous polyposis and an elevated risk of early onset colorectal and duodenal cancer^19–22^. The age of onset and the burden of intestinal polyps are highly variable between individuals, ranging from 10s to 100s leading to a substantially increased incidence of colorectal cancer.^23–27^ Risks of other cancer types are also thought to be increased^28^.

Colorectal adenomas and carcinomas from individuals with MAP show a predominance of C>A mutations consistent with the presence of an elevated mutation rate attributed to defective MUTYH function^29–33^. However, whether there is an increased mutation rate in normal cells from individuals with biallelic germline *MUTYH* mutations is unknown. If present in normal cells, understanding the magnitude of the increase in mutation rate, the tissues and cell types in which it occurs, the proportion of cells which show it, the mutational processes responsible and the effects of early neoplastic change would provide insight into the genesis of the elevated cancer risk observed in these individuals. In this study we characterise the somatic mutation rates and mutational signatures of normal intestinal cells, and other normal cells, from individuals with MAP.

## RESULTS

### Clinical information

Ten individuals aged 16 to 79 years with biallelic germline *MUTYH* mutations were studied. These included five missense mutation homozygotes (four MUTYH^Y179C+/+^, one MUTYH^G286E+/+^), three compound heterozygotes for the same pair of missense mutations (MUTYH^Y179C+/-;G396D+/-^), and two siblings homozygous for a nonsense mutation (MUTYH^Y104*+/+^). These *MUTYH* germline mutations have all been previously recognised as predisposing to MAP^22,23^. All 10 individuals had colorectal polyposis, with between 16 and >100 colonic adenomas, six were known to have duodenal polyps, five had colorectal cancer and one developed jejunal and pancreatic neuroendocrine cancer (Extended Data Table 1).

### Mutation rates in normal intestinal stem cells

An intestinal crypt is constituted predominantly of a population of epithelial cells arising from a single recent common ancestor^34–36^. The somatic mutations which have accumulated over the course of the individual’s lifetime in the ancestral crypt stem cell are present in all its descendant cells^3^. Thus, by sequencing individual crypts, somatic mutations present in the ancestral stem cell can be identified. Using laser-capture microdissection, 144 individual normal intestinal crypts (large intestine n=107 and small intestine n=37) were isolated from the 10 individuals with germline *MUTYH* mutations (Extended Data Table 2). DNA libraries were prepared from individual crypts using a bespoke low-input DNA library preparation method^37^ and were whole-genome sequenced at a mean 28-fold coverage.

The single base substitution (SBS) mutation burdens of individual crypts ranged from a median for each individual of 2,294 to 33,350, equating to mutation rates of 79-1470 SBS/year, 2-33-fold higher than normal crypts from wild-type individuals (~43 SBS/year)^3^ (Figure 1b). Therefore, all normal crypts from all MAP individuals studied showed elevated somatic mutation rates (Figure 1a-b).

**FIGURE 1.**
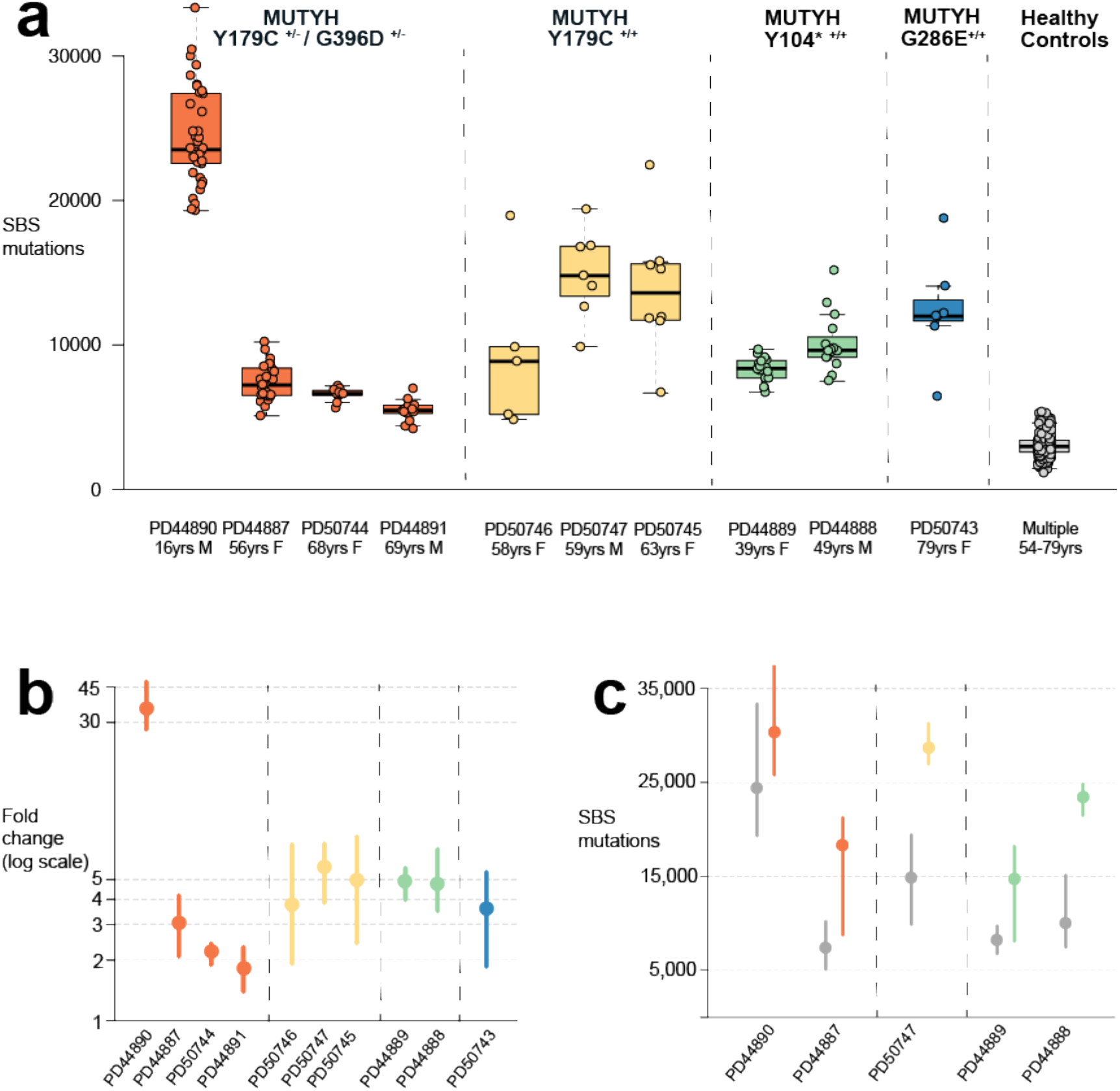
Somatic mutation burdens in cells with *MUTYH* mutations. Elevated mutation burdens in normal intestinal cells with *MUTYH* mutations (a) Genome-wide single base substitution (SBS) mutation burden of individual intestinal crypts grouped according to patient. Each dot represents an individual intestinal crypt. *MUTYH* genotypes are displayed separately. Boxplots display median, inter-quartile range (IQR) from 1^st^ to 3^rd^ quartiles and whiskers extend from the last quartile to the last data point that is within 1.5x IQR (b) Fold-change in SBS rate compared with wild-type controls^3^ coloured according to germline genotype (orange; MUTYH^Y179C+/- G396D+/-^, yellow; MUTYH^Y179C+/+^, green; MUTYH^Y104*+/+^, blue; MUTYH^G286E+/+^). Median values are represented by the dot, whiskers represent the range from the minimum to maximum value. P-values for pair-wise comparisons are shown in Supplementary Note. (c) Genome-wide single base substitution burden in histologically normal crypts (grey) and adenoma crypts (orange, yellow and green) arranged by patient and germline mutation. Median values represented by the dot, whiskers represent the range from the minimum to maximum value. Data available for 5 individuals who had adenoma crypts sequenced.

Differences in mutation rate were observed between individuals with MAP (Figure 1b). A 33-fold higher rate of SBS accumulation than in wild-type crypts^3^ was observed in PD44890, a 16 year old male with MUTYH^Y179C+/- G396D+/-^. By contrast, the nine other individuals showed only 2- to 5-fold increases in mutation rate compared to wild type. The reason for this substantial difference is not clear. However, in addition to the *MUTYH* mutations, PD44890 carried two heterozygous germline missense variants in *OGG1* (Extended Data Figure 1), one of which (R46Q) is reported to impair OGG1 activity in experimental systems^38,39^ and has been observed as somatically mutated in human cancer^40^. Germline *OGG1* mutations are not currently recognised as causing cancer predisposition in humans^41^. However, if either or both of these mutations results in defective 8-OG excision they could account for the substantially elevated mutation rate in PD44890, particularly in the context of defective MUTYH activity.

There was also evidence of differences in mutation rates between the various *MUTYH* germline genotypes studied (Fig. 1b). Excluding the outlier individual PD44890, mutation rates were lower in individuals with the compound heterozygous MUTYH^Y179C+/- G396D+/-^ (102 SBS/year, range 61-182) than individuals with MUTYH^Y179C+/+^ (214 SBS/year, range 84-356), MUTYH^Y104*+/+^ (204 SBS/year, range 153-309) or MUTYH^G286E+/+^ (152 SBS/year, range 82-238) (MUTYH^Y179C+/- G396D+/-^ vs MUTYH^Y179C+/+^ / MUTYH^Y104*+/+^ / MUTYH^G286E+/+^, *P*=9.5×10^-8^, *P*=6.1×10^-19^, *P*=9.8×10^-3^ respectively. MUTYH^G286E+/+^ vs. MUTYH^Y179C+/+^ / MUTYH^Y104*+/+^ *P*=2.7×10^-2^ and *P*=4.5×10^-3^ respectively). The results, therefore, indicate that different *MUTYH* genotypes confer differentially elevated mutation rates and that the extent of the mutation rate increase can be modified by other factors.

SBS mutation rates in coding exons in normal intestinal crypts from MAP individuals were also elevated compared to wild-type individuals (Extended Data Fig 2a-b). These increases were, however, slightly smaller than those observed in the genome-wide mutation rate (Extended Data Fig 2a-b). Nonsense, missense and synonymous mutation rates were all increased compared with wild-type crypts, with the greatest increase observed in nonsense mutations (~10-fold more nonsense than wild-type vs ~3.5-fold more missense and ~2.6-fold more synonymous) (Extended Data Fig. 2c). This is attributable to the mutational signatures present (see below) and the tendency of specific mutations at particular trinucleotide contexts to preferentially generate protein-truncating mutations^42,43^.

Neoplastic glands from 13 intestinal adenomas from five individuals with MAP showed SBS mutation burdens that were, on average, ~2-fold higher (range 1.2 to 2.5 fold) (Figure 1c) than normal crypts from the same individuals sampled at the same time. Therefore, the elevated mutation rate observed in histologically normal intestinal crypts in individuals with germline *MUTYH* mutations is further increased during the process of neoplastic transformation, as previously observed in wild type individuals^44,45^.

Small insertion and deletion (ID) mutations accumulated at a rate of 2.1 ID/yr (linear mixed-effects model, 95% confidence interval (C.I.) 1.2-3.0, *P*= <10^-4^), which is higher than in wild-type controls (1.3 ID/yr, linear mixed-effects model, C.I. 0.54-2.0, *P=* 0.0011)^3,43^. The cause of this modestly elevated ID rate is not clear. In two MAP individuals additional mutational processes could explain the higher burdens observed in these cases. In PD44890 the high ID mutation rate (ID rate 6/yr) was, at least partially, explained by the presence of an additional ID generating mutational process associated with exposure to the mutagen colibactin produced by a strain of *E.Coli*^3,46,47^ present in the colonic microbiome of some people (see below). In PD50747 (ID rate 6/yr), a previously undescribed sporadic ID signature IDA was identified which was not present in other MAP individuals (described below). Structural rearrangements and copy number changes were only observed in a small number of normal intestinal crypts, at similar frequencies to those in wild-type controls (Extended Data Table 2)^3^.

### Mutational signatures

Mutational signatures were extracted from the combined catalogues of SBS mutations from all normal and neoplastic intestinal crypts and glands using two independent methods. We then decomposed each *de novo* extracted signature into known COSMIC reference mutational signatures. Finally, we used these decompositions to estimate the contribution of each reference signature to each sample (Methods, Supplementary Note). Three *de novo* extracted signatures, N1-N3, accounted for the majority of mutations, all of which were mainly characterised by C>A mutations (Fig. 2a). Two of these closely corresponded to the reference mutational signatures SBS18 and SBS36. The third was abundant only in individual PD44890 (the individual with a high mutation rate carrying *OGG1* germline variants) (Fig. 2b and Supplementary Note) but could also be accounted for by a combination of SBS18 and SBS36 according to the standard parameters used for decomposition (Supplementary Information).

**FIGURE 2.**
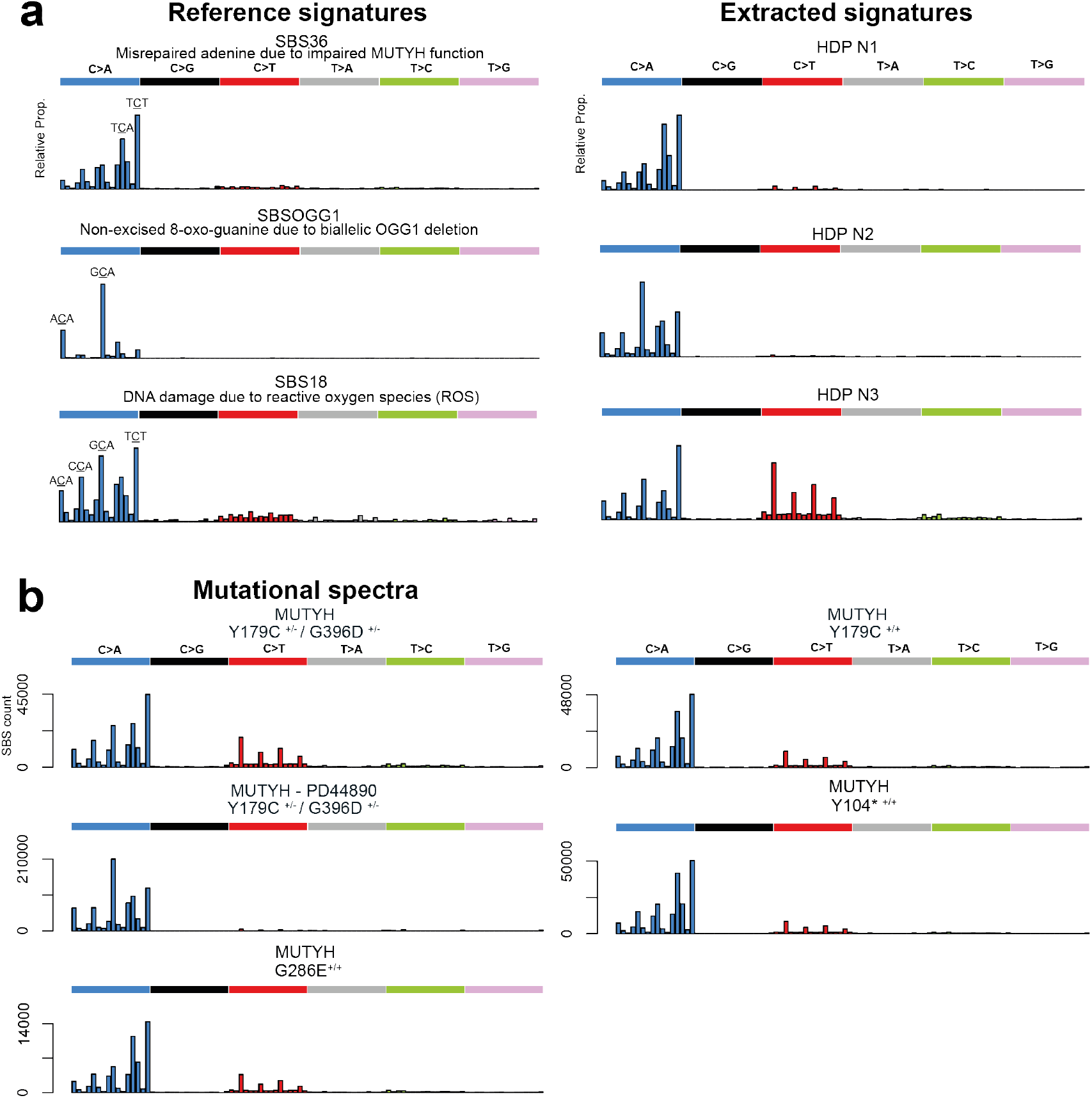
Mutational spectra and signature components from normal cells with MUTYH mutations. **(a)** Probability distribution for COSMIC reference signature SBS36^40^, recently described *OGG1* deletion signature SBSOGG1^51^ and reference signature SBS18^40^. Mutational signature components N1-3 from HDP *de novo* signature extraction (see Supplementary Information for all components and Methods / Supplementary Note for further explanation). **(b)** Mutational spectra in normal tissues displayed by the germline *MUTYH* mutation. Distinctive peaks are annotated with their trinucleotide context (mutated base is underlined). PD44890 is displayed separately to highlight the difference in spectrum observed in this individual.

Following decomposition and signature attribution, four reference SBS mutational signatures, SBS1, SBS5, SBS18 and SBS36 were identified in all samples (Fig. 3, Supplementary Note). SBS1, due to deamination of 5-methlycytosine at CG dinucleotides and SBS5, of unknown aetiology, have both been found ubiquitously in normal and cancer cells and accumulate in a more or less linear fashion with age^2–4,8,33,48–50^. SBS18, thought to result from DNA damage due to reactive oxygen species, has previously been reported in normal colorectal cells^3^ and many types of cancer^33^ and is characterised by C>A mutations predominantly at ACA, CCA, GCA and TCT trinucleotide contexts (mutated base underlined) (Fig. 2a and Fig. 3). SBS36 has previously been found in cancers with germline or somatic *MUTYH* mutations and is also characterised by C>A mutations, albeit with a different profile of preferred trinucleotide contexts from SBS18^30–33^(Fig. 2a). SBS88, which is predominantly characterised by T>C and T>G mutations, and is due to early life exposure to the mutagenic agent colibactin produced by some strains of *E.Coli*^3,46,47^, was observed in a subset of crypts from PD44890 (Figure 3b). The SBS88 mutation burdens were consistent with those previously seen in wild type individuals indicating that MUTYH is unlikely to be implicated in the genesis of SBS88.

**FIGURE 3.**
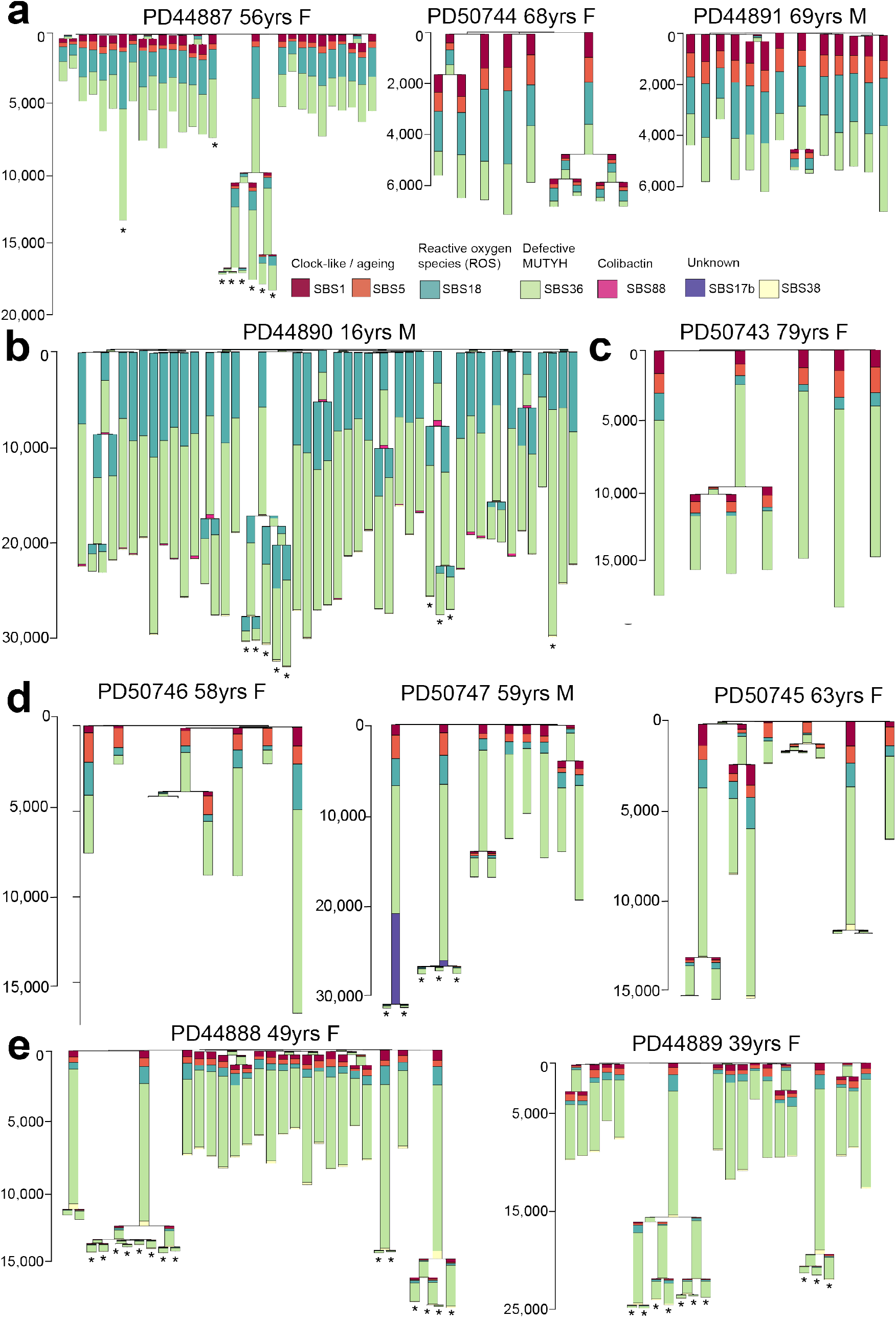
Phylogenetic trees and mutational signatures intestinal cells with germline *MUTYH* mutations. Phylogenetic trees per-individual reconstructed from SBS mutations in individual intestinal crypts showing the number of SBS mutations per branch. Stacked barplots are overlaid onto each branch to represent the proportion of each mutational signature contributing to that branch. Phylogenetic trees are arranged by *MUTYH* germline mutation; (a) MUTYH^Y179C+/- G396D+/-^ (b) MUTYH^Y179C+/- G396D+/-^ with OGG1 germline mutations, (c) MUTYH^G286E+/+^, (d) MUTYH^Y179C+/+^ and (e) MUTYH^Y104*+/+^. Adenoma glands bearing cancer driver mutations are indicated with an asterisk ‘*’.

The increased SBS mutation burdens in normal crypts from individuals with *MUTYH* germline mutations appeared to be due to the contributions of SBS18 and SBS36 mutations (Fig. 3a-e). The proportions of SBS18 and SBS36, however, differed between *MUTYH* germline genotypes. SBS18 accounted for a substantially higher proportion of mutations in crypts and glands from individuals with the MUTYH^Y179C+/- G396D+/-^ genotype (n=85 crypts) than in individuals with the MUTYH^Y179C+/+^, MUTYH^Y104*+/+^ and MUTYH^G286E+/+^ genotypes (n=59 crypts, Extended Data Fig. 3a). Since MUTYH^Y104^* causes MUTYH protein truncation, it is conceivable that SBS36 is the consequence of complete loss of MUTYH function and therefore that this is also effected by MUTYH^Y179C^ and MUTYH^G286E^. Conversely, MUTYH^G396D^ may retain partial activity^14,21^ and thus generates a signature more closely resembling SBS18 which is found in normal tissues with fully active MUTYH.

The *de novo* extracted mutational signature N2, which primarily contributes to the mutational spectra of crypts from PD44890 (MUTYH^Y179C+/- G396D+/-^), resembled reference signature SBS18 (https://cancer.sanger.ac.uk/cosmic/signatures) but showed differences, notably with over representation of C>A mutations at GCA and, to a lesser extent, CCA and ACA trinucleotides (mutated base underlined) (Fig. 2a-b and Extended Data Fig. 1). A signature reported in human cells with *in vitro* engineered biallelic *OGG1* deletion is also primarily characterised by C>A mutations at GCA and ACA trinucleotides^51^. It is, therefore, possible that mutagenesis due to the germline *OGG1* variant(s) in PD44890 (see above) is superimposed on the mutational signature produced by the *MUTYH* germline mutations to generate N2 (see Supplementary Information for further analysis and discussion).

The mutational signatures in adenoma glands were similar to those seen in normal crypts from the same individuals (Fig. 3a,b,d,e). SBS36 and SBS18 were principally responsible for the increased mutation burdens observed in adenomas compared to normal crypts.

Candidate cancer “driver” mutations, defined as known or likely oncogenic hotspot mutations and truncating mutations in tumour suppressor genes (Methods), were observed in 15% of normal crypts (22/144), more than double the rate observed in wild-type crypts from comparable healthy controls; 6% (25/449)^3,43^. A substantial proportion of candidate drivers (16/22) were nonsense mutations, mirroring the broader exome-wide increase in nonsense mutations (Extended Data Fig. 2c), and reflecting the proclivity of certain mutation types to generate truncating mutations^29,42,43^. The mutational spectrum of driver mutations in normal crypts and neoplastic glands resembled the genome-wide spectra with substantial contributions from SBS18 and SBS36 (Extended Data Fig. 4). Hence, the mutational processes resulting from defective MUTYH activity appear to promote the accumulation of putative cancer driver mutations in normal and neoplastic tissues^52,53^.

Three known ID signatures were identified. ID1 & ID2 are characterised predominantly by insertions and deletions of single T bases at T mononucleotide repeats which are associated with strand slippage during DNA replication and are seen in most human cancers and normal tissues^1–4,8,33^. ID18 is associated with colibactin exposure, is found in normal intestinal stem cells and certain cancers, and usually associated with SBS88^3,47^. ID1 was the dominant signature in normal cells whereas ID2 predominated in neoplastic cells (Extended Data Fig. 5). ID18 was principally observed in samples from PD44890 and is responsible for the elevated ID rate in this individual (Extended Data Fig. 6). A further ID signature, IDA, identified in PD50747, was characterised by single C insertions at C mononucleotide repeats (Extended Data Fig. 6). IDA was present in both normal crypts (~5% of total ID burden) and to a greater extent in adenoma glands (~20% of total ID burden). The cause of this previously undescribed signature is unclear but may be associated with previous capecitabine treatment and seems unlikely to be related to germline MUTYH mutations.

### Mutations in other cell types

To investigate whether the elevated mutation rates and mutational signatures observed in intestinal epithelium caused by defective MUTYH are present in other cell types, peripheral white blood cell and tissue lymphocyte DNAs from individuals with biallelic *MUTYH* mutations were whole genome sequenced using a duplex sequencing method (NanoSeq)^50^ that allows mutation calling from single DNA molecules and thus accurately discovers somatic mutations in tissues in which multiple clonal lineages are intimately mixed.

The white blood cell SBS mutation rates of all individuals with *MUTYH* mutations were higher than wild-type controls (*n*=15 granulocyte samples from 9 healthy individuals aged 20-80yrs)(Fig. 4)(25 SBS/yr vs 19 SBS/yr, linear mixed-effects model, *R*^2^=0.89, MUTYH; 95% C.I., 19-31, *P*=10^-7^ and wild-type; 95% C.I., 14-24, *P*=10^-6^). The relative increases in blood mutation rates were lower than in intestinal crypts from each individual (Fig. 4b). Nevertheless, the relative increases paralleled the differential increases observed between individuals in intestinal crypts. SBS mutation rates in tissue lymphocytes were modestly raised compared with wild-type healthy individuals (Fig. 4d) (53 SBS/yr vs 40 SBS/yr, linear mixed-effects model, *R*^2^=0.68, *MUTYH;* 95% C.I., 21-85, *P*=0.01 and wild-type; 95% C.I., 13-66, *P*=0.01). The signatures associated with defective MUTYH, SBS18 and SBS36, contributed the excess mutations in all samples (Fig. 4c and Fig. 4e). An additional mutational signature was seen in lymphocytes. SBS9, which is associated with DNA polymerase eta mediated somatic hypermutation and is a key process in the physiological maturation of B-cells, was observed in most lymphocyte samples indicating that the lymphocyte cell populations contained mature B-cells (Fig. 4e).

**FIGURE 4.**
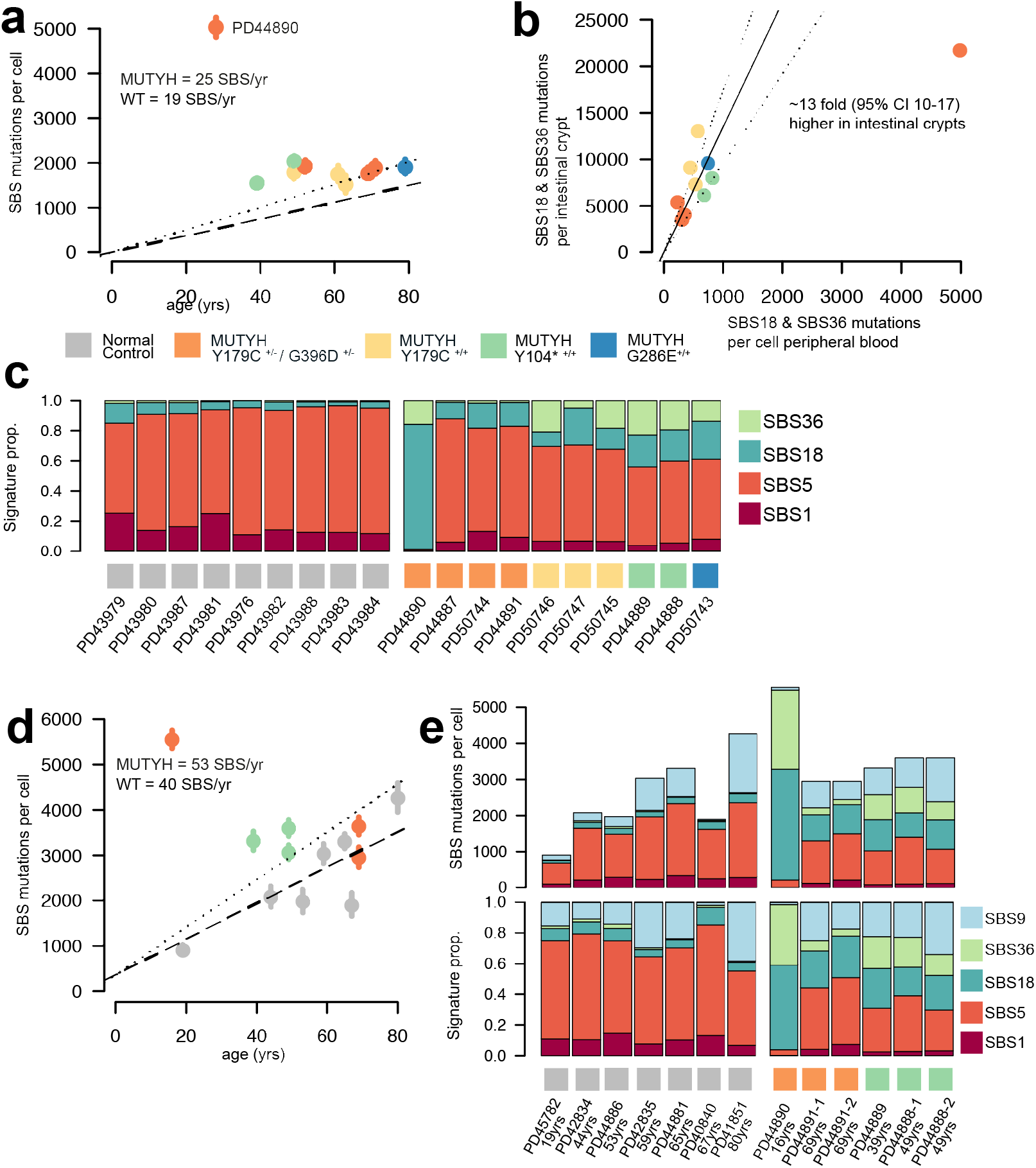
Mutation burdens and mutational signatures in blood and immune cell populations. **(a)** Mutation burden in peripheral blood. SBS mutation burden per cell plotted against the age of the individual in years coloured according to the individual’s germline mutation; orange; MUTYH^Y179C+/- G396D+/-^, yellow; MUTYH^Y179C+/+^, green; MUTYH^Y104*+/+^ and blue; MUTYH^G286E+/+^. Whiskers represent the 95% confidence interval. Black dashed line represents the average mutation rate in normal wild-type control samples^50^ **(b)** Mutation burden of *MUTYH* associated mutational signatures; SBS18 and SBS36 per cell for peripheral blood (x-axis) against the SBS18 & SBS36 mutation burden of normal intestinal crypts (y-axis). Each dot represents one individual and they are coloured according to the individual’s germline mutation; orange; MUTYH^Y179C+/- G396D+/-^, yellow; MUTYH^Y179C+/+^, green; MUTYH^Y104*+/+^ and blue; MUTYH^G286E+/+^. Ratio MUTYH mutational processes in colon vs blood ~13x fold (linear model, 95% C.I.; 10-17). Black line indicates ratio and dotted lines the 95% confidence intervals. **(c)** Stacked bar plot displaying the mutational signature contribution in each peripheral blood sample organised by patient. Coloured squares indicate the *MUTYH* germline mutation. Normal control data from granulocytes sequenced using the same method (data from Abascal et al 2021)^50^. Significantly higher proportion of SBS18 and SBS36 are observed in individuals with MUTYH mutations vs normal healthy controls (two-sided Wilcoxon rank sum, *P*=0.00004) **(d)** SBS mutation burden in intestinal lymphocyte cells from wild-type healthy individuals (grey) and individuals with *MUTYH* mutations (coloured according to the germline *MUTYH* genotype) plotted against age (years). Dots represent median values and whiskers represent the 95% confidence interval. Dashed line indicates the rate of increase of SBS burden in wild-type lymphocytes (40 SBS/yr, linear mixed-effects model, *R*^2^=0.68, 95% C.I., 13-66, *P*=0.01) and dotted line indicates the rate of increase in SBS burden in lymphocytes from individuals with *MUTYH* mutations (53 SBS/yr, linear mixed-effects model, *R*^2^=0.68, 95% C.I., 21-85, *P*=0.01). **(e)** Stacked bar plots showing the absolute (above) and relative (below) contributions of each mutational signature in tissue lymphocytes from wild-type healthy individuals (grey squares) and individuals with *MUTYH* mutations (coloured squares).

## DISCUSSION

This study shows elevated base substitution somatic mutation rates due to SBS18 and/or SBS36 in normal tissues from individuals with *MUTYH* mutations. The results are compatible with all intestinal, and potentially all other cells in the body, showing elevated mutation rates. The relative increases in mutation rate and mutational signature composition differed between individuals, probably due to different *MUTYH* mutations and perhaps to other modifying influences.

We have previously highlighted the capability of normal human cells to tolerate substantially elevated mutation rates^43^. Carriers of POLE and POLD1 germline exonuclease domain mutations exhibited elevated somatic mutation burdens without evident cellular or organismal consequences, other than an increased cancer risk^43^. This capability is confirmed in *MUTYH* germline mutation carriers. It is further emphasised by the observation of a 33-fold genome-wide elevated base substitution mutation burden in the 16 year old PD44890, which would confer a “mutational age” of ~500 years, without overt evidence of premature ageing. The increase in mutation burden in coding exons is lower than genome-wide in POLE/POLD1 mutation carriers. Similarly, in individuals with *MUTYH* mutations there is a smaller increase of coding exon than genome-wide mutation burdens (Extended Data Fig. 2a-b). Nevertheless in PD44890 the increase is still ~29-fold, and therefore equivalent to a “mutational age” of ~450 years. Whilst lesser increases in mutation rates compared to wild-type individuals were observed in other tissues from PD44890, ~8-fold in white blood cells and ~7 fold in tissue lymphocytes, these still conferred substantially elevated “mutational ages” in the absence of features of premature ageing. Thus direct deleterious effects of base substitutions accumulated over the course of a lifetime may not be an important cause of ageing.

The elevated mutation rate in normal intestinal epithelium likely contributes to the increased risk of colorectal adenomas and cancers in individuals with *MUTYH* mutations. Indeed, there appears to be a correlation between the extent of elevation of mutation rate and the rate of acquisition of colorectal adenomas. Individuals with the MUTYH^Y104*+/+^ and MUTYH^Y179C+/+^ genotypes exhibited greater increases in somatic mutation rates than individuals with the MUTYH^Y179C+/- G396D+/-^ genotype. Previous detailed clinical phenotyping of large series indicates that individuals with biallelic truncating mutations or MUTYH^Y179C+/+^ show higher rates of accumulation of adenomas and earlier age of onset of carcinoma^22,23^ than MUTYH^Y179C+/- G396D+/-^. The correlation between elevation of mutation rate and severity of clinical phenotype is further highlighted by individual PD44890 (16 years of age, MUTYH^Y179C+/- G396D+/-^) who exhibited a substantially higher mutation rate than others of this genotype, and showed a much accelerated rate of colorectal adenoma development (Extended Data Table 1-2). We previously described ~7-fold elevated genome-wide base substitution mutation rates in intestinal cells of POLE germline exonuclease domain mutation carriers^43^. POLE mutation carriers, however, show lower colorectal adenoma rates than *MUTYH* biallelic mutation carriers who generally only show 2-5 fold increased mutation rates. This apparent discrepancy may, however, be explained by the genomic distribution of mutations. In POLE mutation carriers there is relative sparing of coding sequences, with only a three to four fold increase in exonic mutations in intestinal cells, whereas this sparing is less pronounced in *MUTYH* mutation carriers leading to similar increases in exonic mutation rates (Extended Data Fig. 7a-b). These observations lead to the proposition that measurements of somatic coding mutation rates undertaken early in life could, in future, be used to refine individual cancer risk predictions for *POLE/POLD* and *MUTYH* germline mutation carriers.

As for many other cancer predisposition syndromes, it is unclear why *MUTYH* germline mutations lead particularly to intestinal neoplasia. Elevated somatic mutation rates are also found in white blood cells in MAP individuals (and may therefore be present in other tissues) although the increases appear lesser in extent than in intestinal cells. The propensity to generate SBS18 mutations appears greater in wild type intestinal cells than in other cell types^49^ and this may also be contributory.

In summary, we report elevated somatic base substitution rates characterised by distinctive mutational signatures in normal tissues from individuals with MAP. These findings underscore previous observations that elevated somatic base substitution rates are largely tolerated by cells and do not overtly accelerate the process of ageing. It is likely, however, that increased mutation rates in normal intestinal cells throughout life lead to increased rates of accumulation of driver mutations and, hence, the procession of neoplastic clones culminating in cancer.

## EXTENDED FIGURES

**Extended Data Figure 1.**
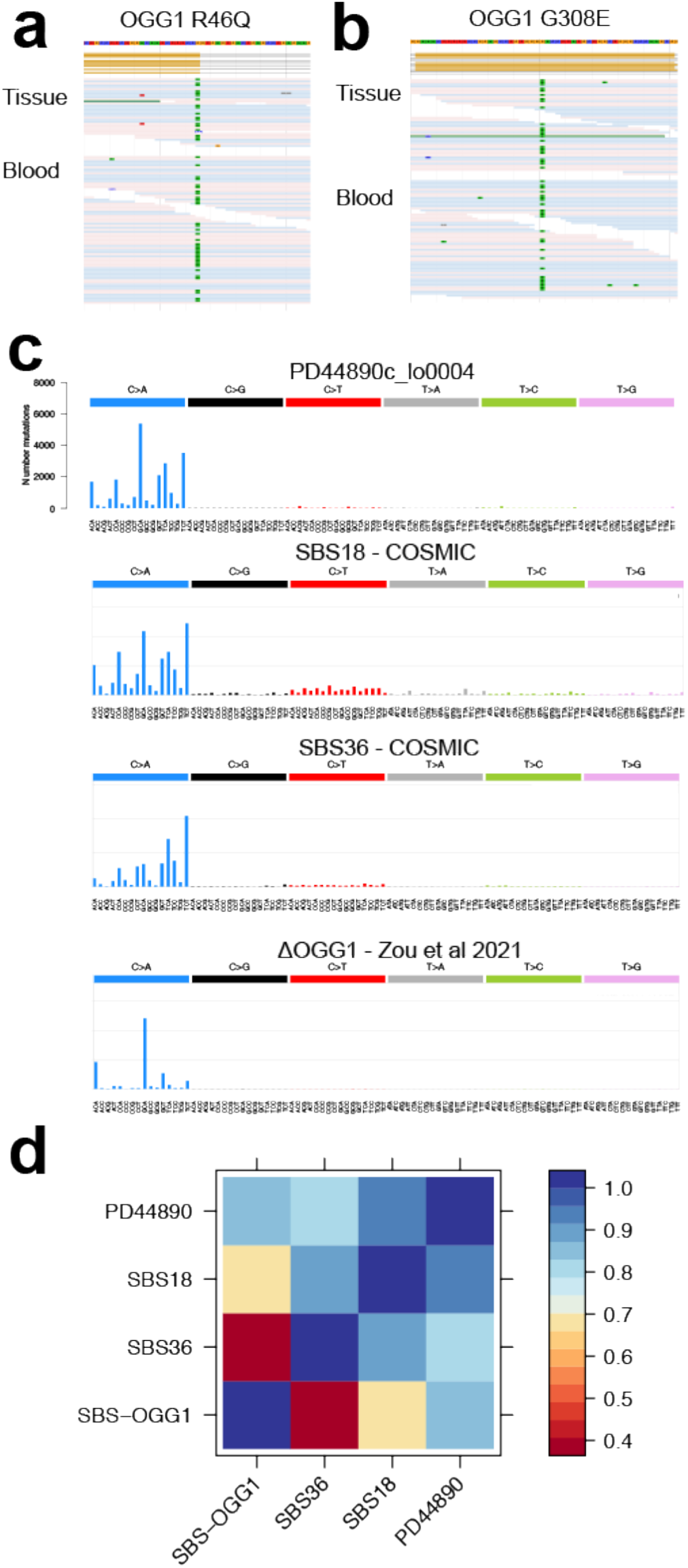
Germline mutations and mutational spectrum of somatic mutations in PD44890. Two candidate germline modifier mutations were identified in individual PD44890 both of which are in the oxoguanine glycosylase (OGG1) gene which is a key component of the base excision repair pathway and a partner of MUTYH in the repair of 8-oxo-guanine DNA lesions. Both mutations are heterozygous nonsynonymous mutations. JBrowse images are displayed showing the reads for each mutation; **(a)** OGG1 R46Q **(b)** OGG1 R308E. See Supplementary Note for further discussion of these putative germline modifier mutations. **(c)** Mutational spectrum of SBS mutations from a representative intestinal crypt from individual PD44890, mutational spectra for the COSMIC reference signatures SBS36 and SBS18 are displayed below for comparison (https://cancer.sanger.ac.uk/cosmic/signatures). Mutational spectrum of IPS cells with homozygous OGG1 deletion (Zou et al 2021, Nature Cancer)^51^ denoted SBS OGG1. **(d)** Matrix displaying cosine similarity between the observed spectrum in individual PD44890 and reference signatures SSB18, SBS36 and SBS OGG1.

**Extended Data Figure 2.**
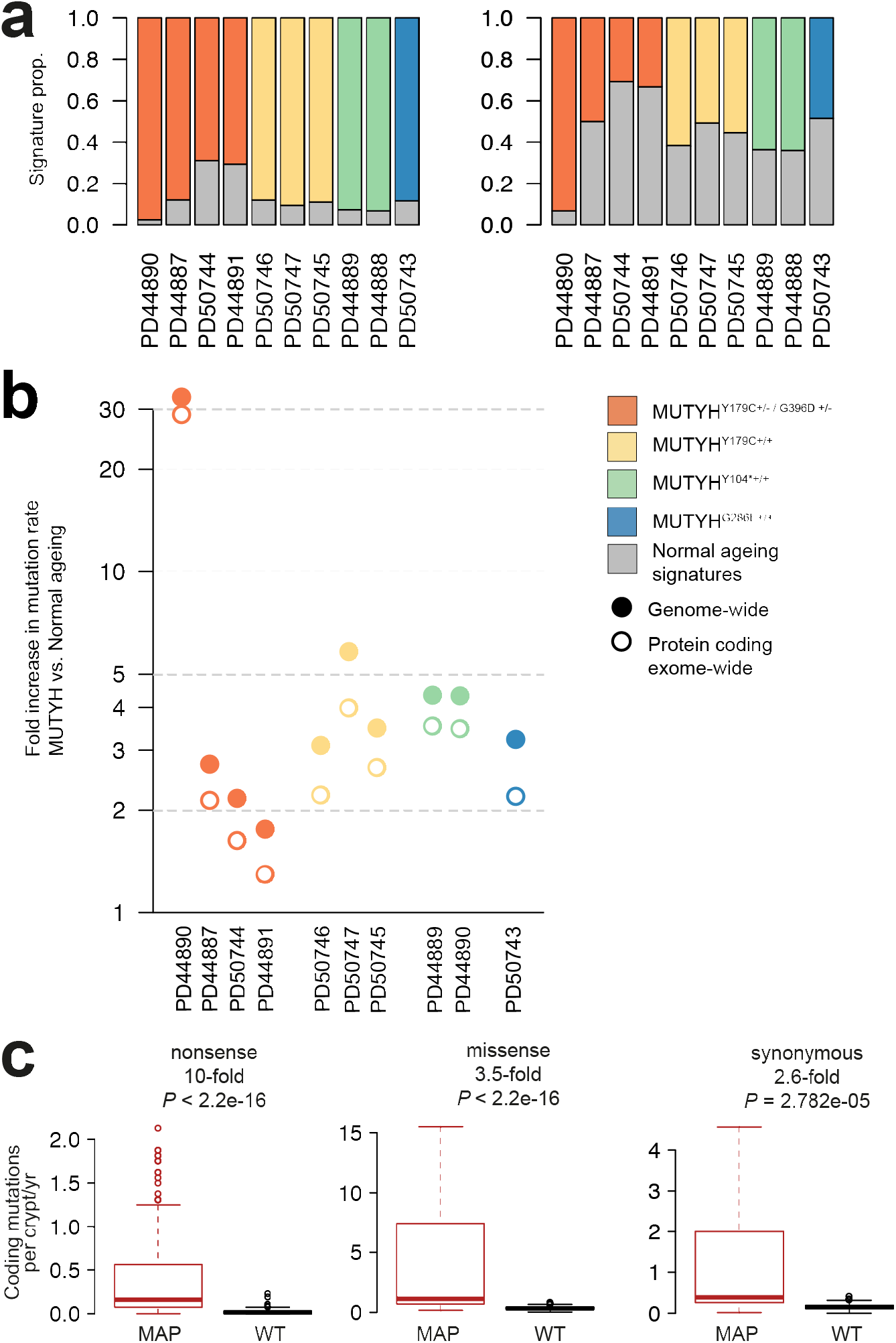
Genome-wide and exome-wide mutation burdens in intestinal crypts. **(a)** Stacked bar plots show the proportion of SBS mutations contributed by MUTYH-related signatures SBS18 and SBS36 (coloured bar) and normal ageing signatures (SBS1 and SBS5) (grey bar). Genome-wide signature proportions (left) protein-coding exome-wide signature proportions (right). **(b)** Fold-increase in the genome- and exome-wide mutation burden of samples from individuals with MUTYH-Associated Polyposis (MAP) compared with normal controls plotted on a logarithmic scale. **(c)** Boxplots showing nonsense, missense and synonymous coding mutation rates (SBS mutations/crypt/year) in histologically normal intestinal crypts from individuals with MUTYH-Associated Polyposis (MAP) (red) and healthy individuals; wild type (WT) (black). Boxplots display median, inter-quartile range (IQR) from 1^st^ to 3^rd^ quartiles and whiskers extend from the last quartile to the last data point that is within 1.5x IQR. Fold changes compared to WT, are shown above each pair, *P*-values calculated using a two-sided Wilcoxon test. Data for healthy wild-type individuals from Lee-Six et al 2019^3^

**Extended Data Figure 3.**
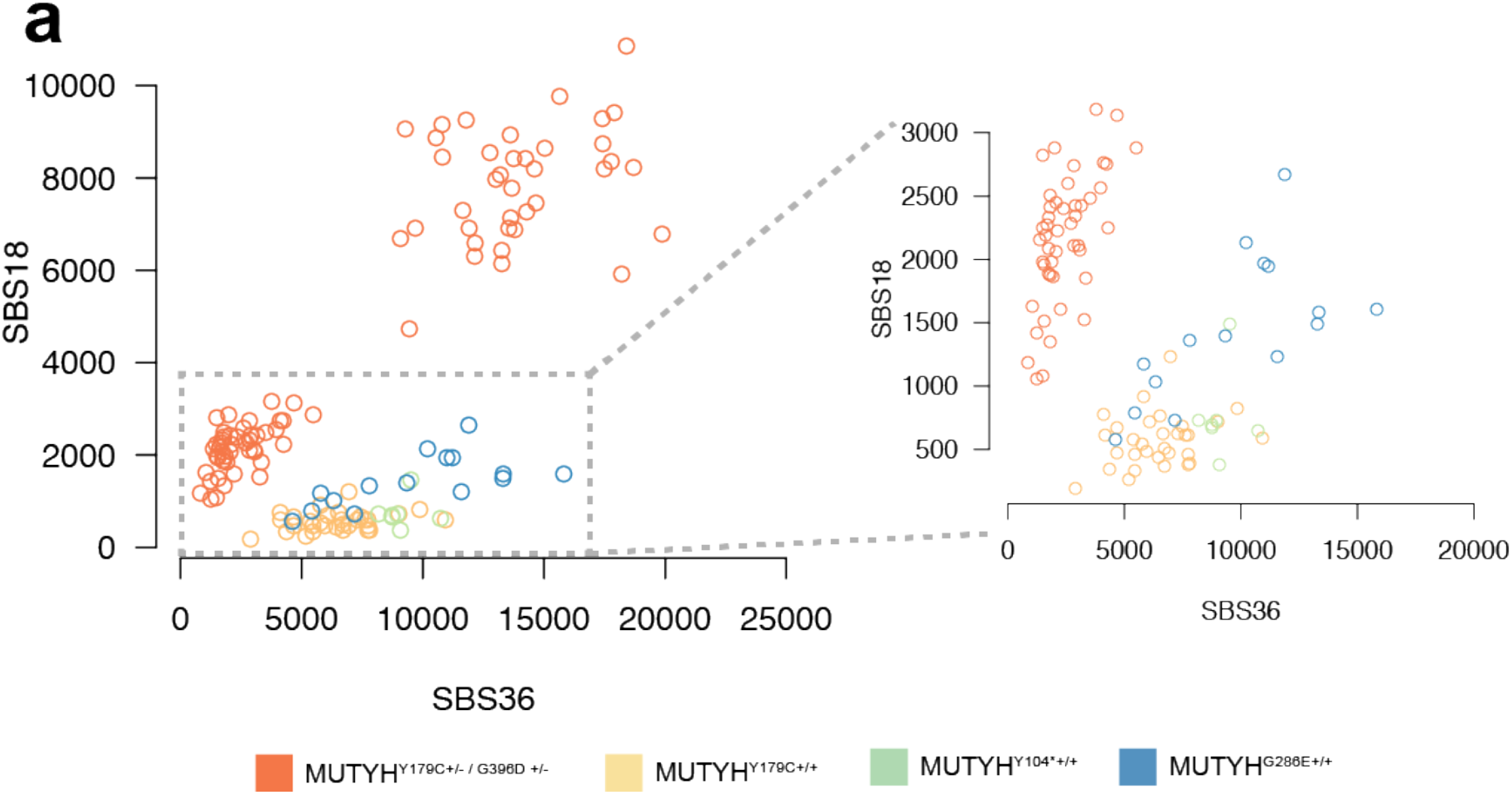
Mutational signatures in normal intestinal crypts and adenoma glands. **(a)** Somatic SBS mutation burden in normal intestinal crypts due to SBS36 (x-axis) and SBS18 (y-axis) coloured according to germline mutation (orange; MUTYH^Y179C+/- G396D+/-^, yellow; MUTYH^Y179C+/+^, green; MUTYH^Y104*+/+^, blue; MUTYH^G286E+/+^). Crypts with MUTYH^Y179C+/+^, MUTYH^Y104*+/+^ and MUTYH^G286E+/+^ show greater relative increases in SBS36 whereas those with MUTYH^Y179C+/- G396D+/-^ show greater increases in SBS18. Crypts from individual PD44890 are a clear outlier with much greater SBS18 burdens. Magnified plot inset.

**Extended Data Figure 4.**
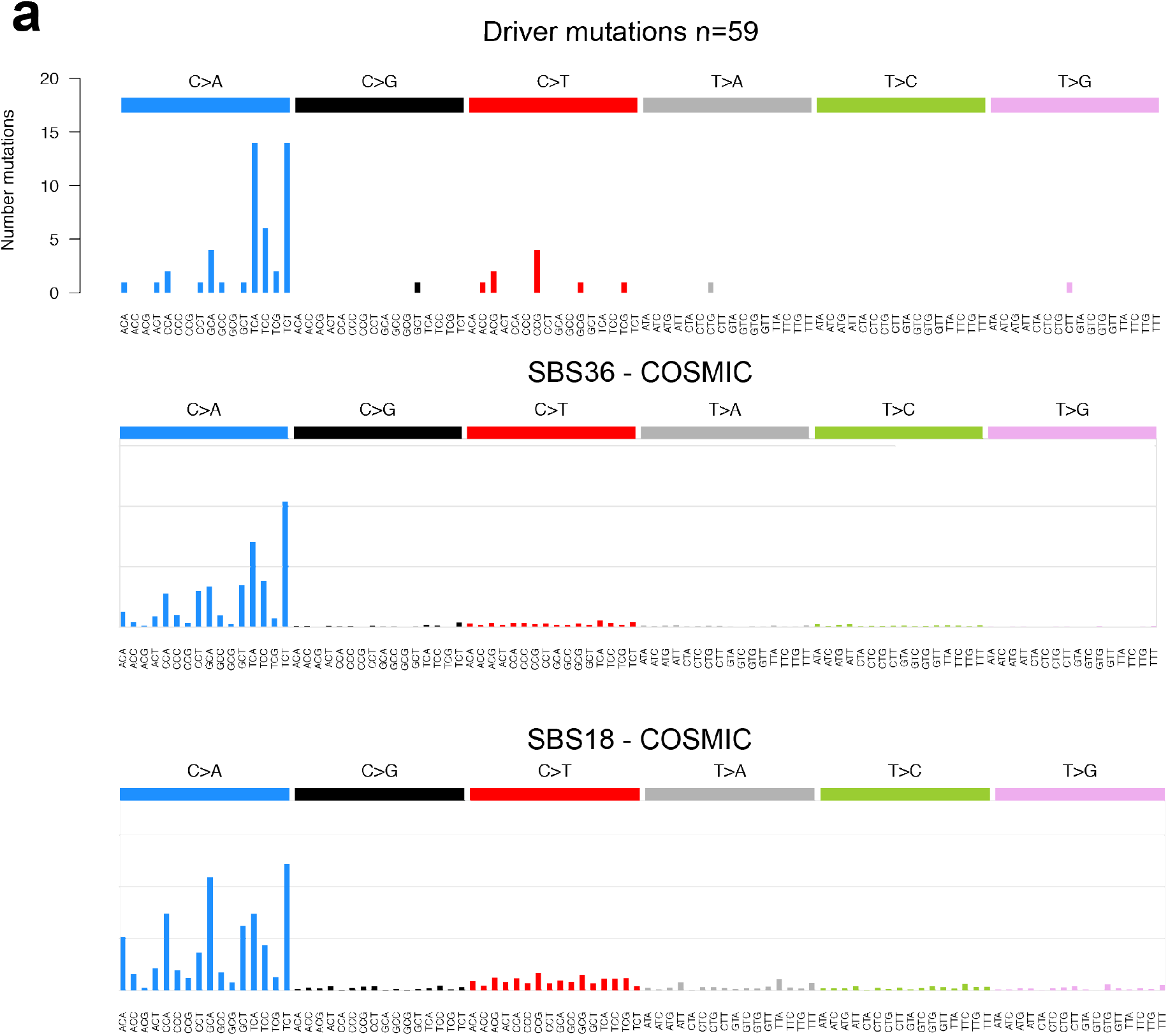
Mutational Spectrum of driver mutations in normal and neoplastic intestinal crypts. **(a)** Mutational spectrum of drivers mutations identified in normal and neoplastic intestinal stem cells in this cohort (n=59). Mutational spectra for the COSMIC reference signatures **(b)** SBS36 and **(c)** SBS18 are displayed below for comparison. The spectra of driver mutations is dominated by SBS36 and SBS18. COSMIC reference signatures available at: https://cancer.sanger.ac.uk/cosmic/signatures.

**Extended Data Fig 5.**
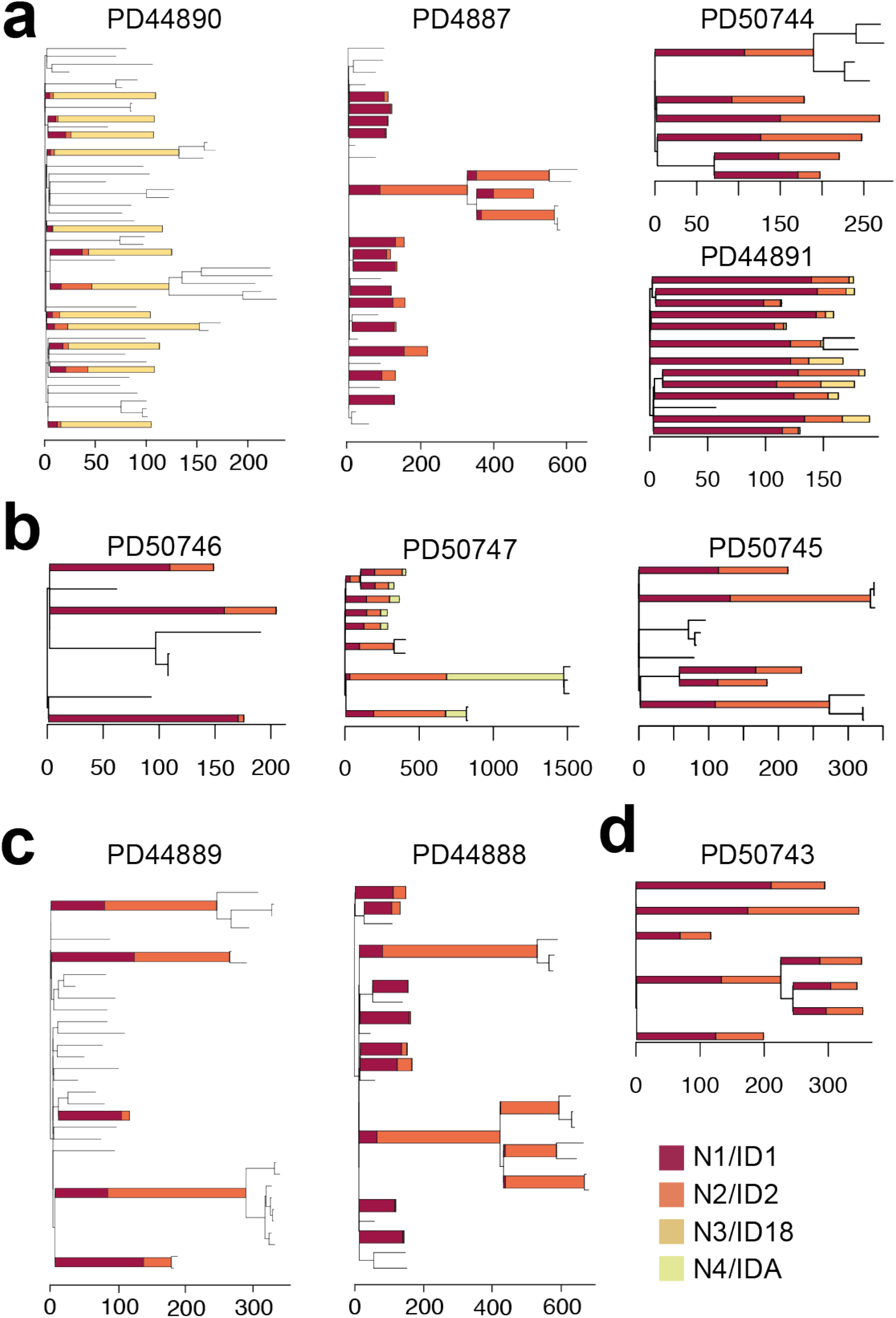
Phylogenetic trees showing ID signature exposure. Phylogenetic trees with ID signature exposure overlaid. Branches of the phylogenetic tree with >100 somatic ID mutations have been converted into stacked bar plots displaying the contribution of each HDP signature component / mutational process to that branch: N1/ID1 shown in dark red, N2/ID2 in orange, N3/ID18 in mustard yellow and N4/IDA in lime green. Phylogenetic tress organised by germline genotype, **(a)** MUTYH^Y179C+/- G396D+/-^, **(b)** MUTYH^Y179C+/+^, **(c)** MUTYH^Y104*+/+^ and **(d)** MUTYH^G286E+/+^

**Extended Data Figure 6.**
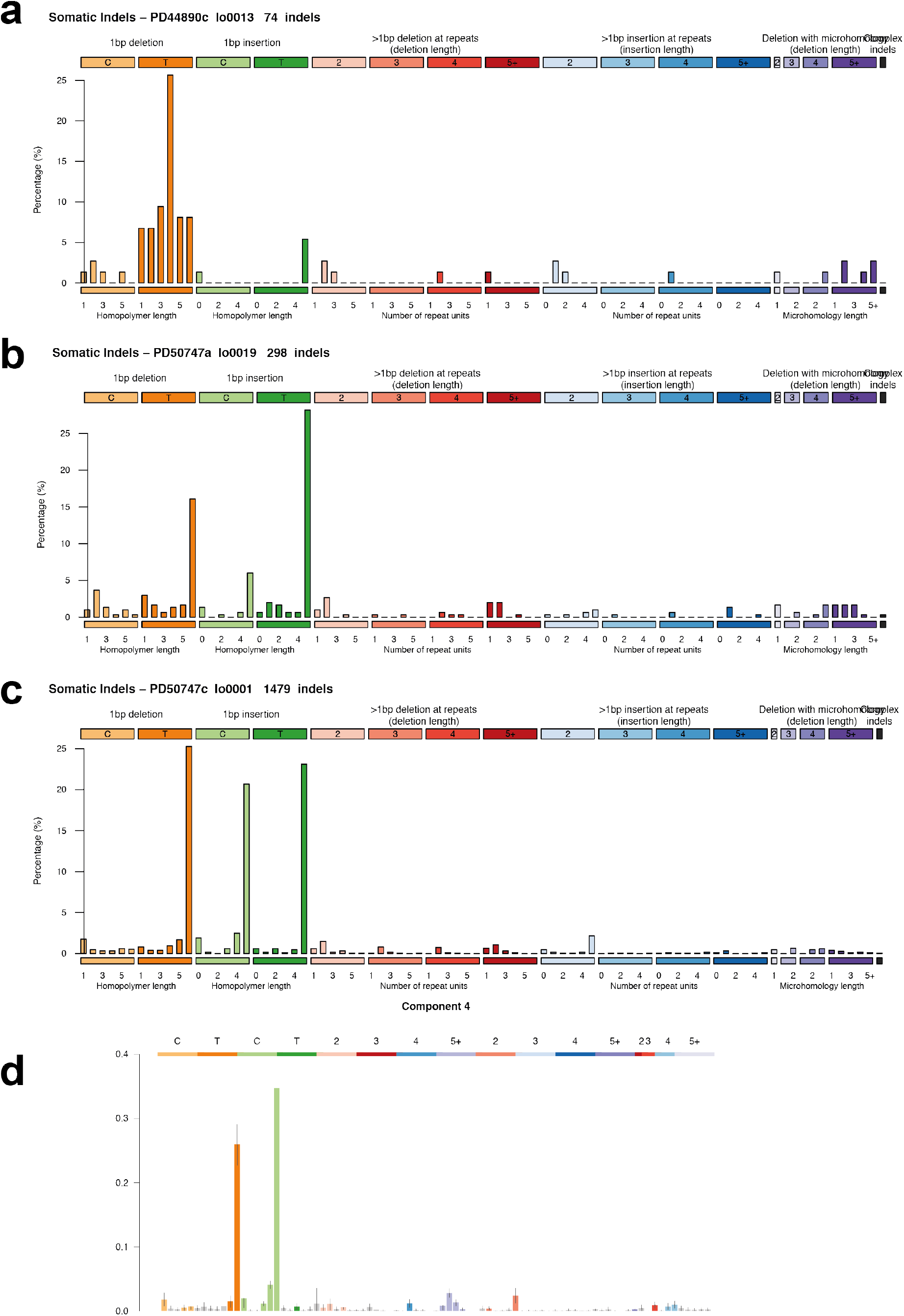
ID mutational spectra. ID mutational spectra for crypts with sporadic ID signature exposures. **(a)** Representative sample from PD44890 with high ID18 exposure. Representative normal crypt from PD50747 showing IDA exposure in **(b)** normal crypt and **(c)** adenoma gland. **(d)** ID signature component arising from unconditioned *de novo* HDP signature extraction (Methods) corresponding to signature IDA.

**Extended Data Figure 7.**
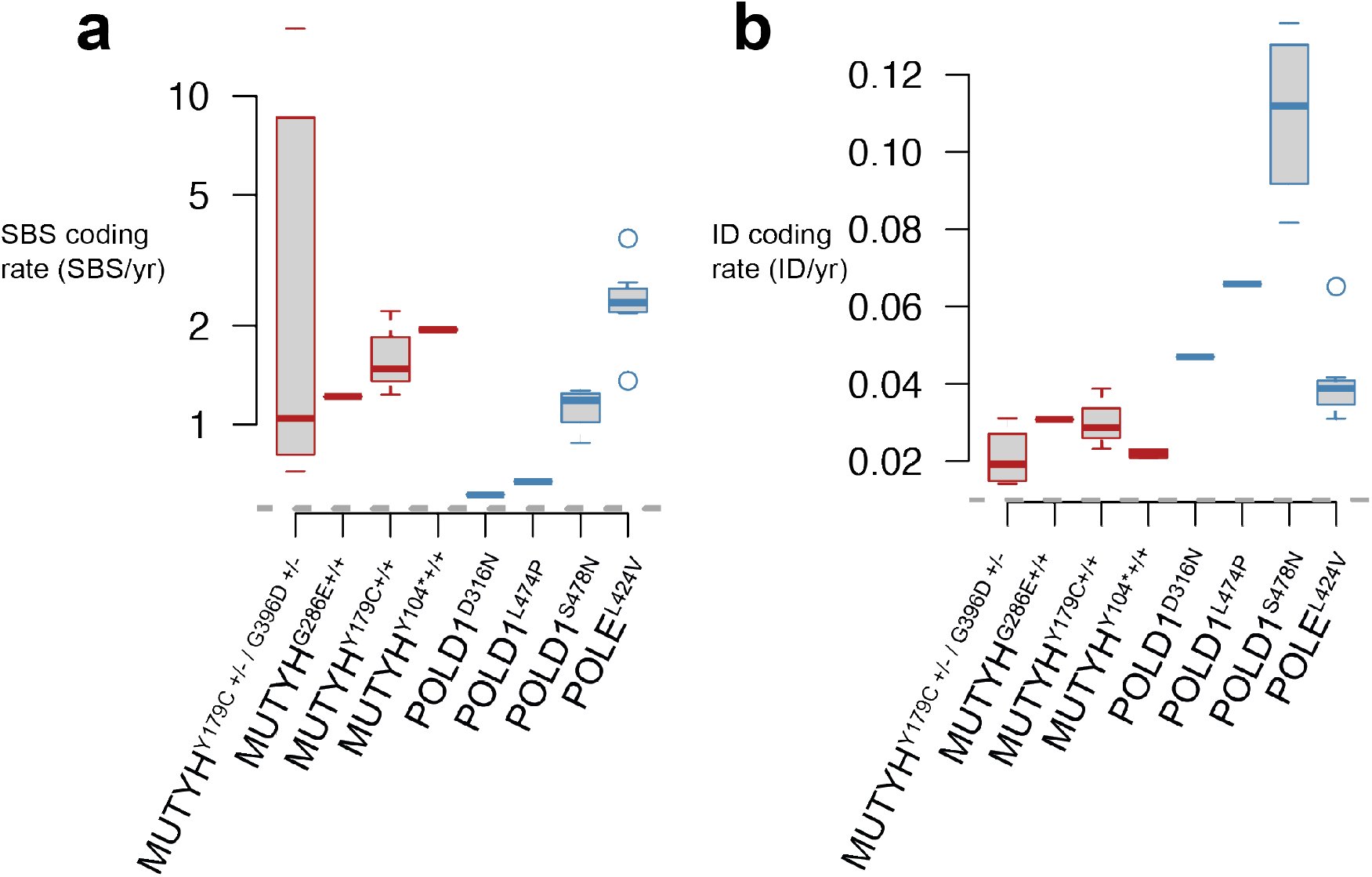
Coding mutation burdens in intestinal crypts from individuals with germline MUTYH and POLE/D1 mutations. Coding mutation burdens in individuals with MUTYH and POLE/D1. **(a)** SBS coding mutation rate (SBS/year) coloured according to cohort; MUTYH; red, POLE/D1; blue. (b) ID coding mutation rate (ID/year).

**Extended Data Table 1 | Clinical summary and phenotypic characteristics of individuals with germline *MUTYH* mutations** Summary of phenotypic features and disease burden in all individuals in this cohort.

**Extended Data Table 2 | Summary of samples sequenced including germline *MUTYH* mutation, sequencing method and mutation burdens**

**Extended Data Table 3 | Cancer driver mutations identified in this cohort** Cancer driver mutations identified across all samples in this cohort

## METHODS

### Ethical approval and study participants

This research complies with all relevant ethical regulations. MAP patients were recruited as part of Wales Research Ethics Committee (REC) 12-WA0071 and 15-WA0075 and samples collected were approved for use in this project by REC 18/ES/0133. Normal healthy controls were recruited as part of the following UK Research Ethics Committee (REC) studies; 15/WA/0131, 15/EE/0152, 18/ES/0133 and 08/h0304/85+5.

Informed consent was obtained from all participants and no monetary compensation was offered for their participation. A complete list of study participants and tissue samples is summarised in Extended Data Table 1 and 2.

### DNA extraction from bulk samples

Frozen whole blood underwent DNA extraction using the Gentra Puregene Blood Kit (Qiagen). Briefly, 1-2ml of frozen blood were thawed, lysed in RBC lysis solution and centrifuged. Cell pellet was resuspended in cell lysis solution and incubated at 37 °C for 2 hours. RNA and protein was degraded using RNase A solution and protein precipitation solution. DNA was precipitated with isopropanol.

### Tissue Preparation

Tissues were embedded in Optimal Cutting Temperature (OCT) compound, frozen histological sections were cut at 25-30μm and mounted on polyethylene naphthalate (PEN) slides and fixed in 70% ethanol for 5 minutes followed by two washes with phosphate buffered saline for 1 minute each. Slides were manually stained in haematoxylin and eosin using a conventional staining protocol. A subset of samples were fixed in RNAlater (Sigma Aldrich) according to manufacturer’s instructions. Fixed tissue samples were embedded in paraffin using a Tissue-Tek tissue processing machine (Sakura). No formalin was used in the preparation, storage, fixation or processing of samples. Processed tissue blocks were embedded in paraffin wax, sectioned to 10μm thickness and mounted onto PEN slides (Leica). Tissue slides were stained using a standard haematoxylin and eosin (H&E) protocol. Slides were temporarily cover-slipped and scanned on a NanoZoomer S60 Slide Scanner (Hamamatsu), images were viewed with NDP.View2 software (Hamamatsu).

### Laser Capture Microdissection

Laser capture microdissection was undertaken using a LMD7000 microscope (Leica) into a skirted 96-well PCR plate. Cell lysis was undertaken using 20μl proteinase-K PicoPure® DNA Extraction kit (Arcturus®), samples were incubated at 65 °C for 3 hours followed by proteinase denaturation at 75 °C for 30 minutes. Thereafter samples were stored at −20 °C prior to DNA library preparation.

### Low-input DNA library preparation and sequencing

DNA library preparation of micro-dissected tissue samples was undertaken as previously described using a bespoke low-input enzymatic-fragmentation-based library preparation method^2,4,37^. This method was employed as it allows for high quality DNA library preparation from very low starting quantity of material (from 100-500 cells). DNA library concentration was assessed after library preparation and used to guide choice of samples to take forward to DNA sequencing, minimum library concentration was 5ng/μL and libraries with >15ng/μL were preferentially chosen. 150bp paired-end Illumina reads were prepared with Unique Dual Index barcodes (Illumina).

DNA sequencing was undertaken on a NovaSeq 6000 platform using an XP kit (Illumina). Samples were multiplexed in pools of 6-24 samples. Pools were sequenced to achieve a coverage of ~30x.

### Mutation calling and post-processing filters

Sequencing reads were aligned to NCBI human genome GRCh37 and aligned using the Burrow-Wheeler Alignment (BWA-MEM). Single Base Substitutions (SBS) were called using the ‘Cancer Variants through Expectation Maximization’ algorithm (CaVEMan)^54^. Mutations were called using an unmatched normal synthetic bam file to retain early embryonic and somatic mutations. Post-processing filters were applied to remove low-input library preparation specific artefacts and germline mutations using a previously described method^1,2,37,55^. Filters applied were: (1) common single nucleotide polymorphisms were removed by filtering against a panel of 75 unmatched normal samples^56^ (2) to remove mapping artefacts, mutations were required to have a minimum median read alignment score of mutant reads (ASMD ≥ 140) and fewer than half of the reads supporting the mutation should be clipped (CLPM =0); (3) a filter to remove overlapping reads that result from the relatively short insert size which could lead to double counting of variant reads; and (4) a filter to remove cruciform DNA structures that can arise during the low-input library preparation method.

Next, we applied multiple filters to remove germline variants and potential artefacts whilst retaining *bona fide* embryonic and somatic variants. This approach has been detailed in previous publications and the code for these filters can be found at https://github.com/TimCoorens/Unmatched_NormSeq. Mutations were aggregated per patient and a read pile-up was performed using an in-house algorithm (cgpVAF) to tabulate the read count of mutant and reference reads per sample for each mutation locus. Germline mutations were filtered out using an exact binomial test. The exact binomial test is used to distinguish germline from somatic variants and uses the aggregate read counts from all samples of the same patient^1,55^. In brief, the read depth across all samples from that individual was calculated (median in this study 496-fold). This high coverage yields a very precise estimate of the true VAF of each mutation. While the VAF estimates of the earliest embryonic SNVs and germline variants from samples sequenced at 30x might overlap, the VAFs from the aggregate coverage from that individual will be distinguishable using statistical testing. To achieve this, the beta-binomial test was applied. The overdispersion parameter (rho) threshold for genuine variants of rho>0.1 was used.

Phylogenetic trees were created using MPBoot (version 1.1.0 bootstrapped - 1000) and mutations were mapped to branches using maximum likelihood assignment.

Indels (ID) were called using Pindel^57^ using the same synthetic unmatched normal sample employed in SBS mutation calling. ID calls were filtered to remove calls with a quality score of <300 (‘Qual’; sum of mapping qualities of the supporting reads) and a read depth of less than 15. Thereafter, ID filtering was performed in a similar manner as SBS to remove germline variants and library preparation / sequencing artefacts.

### Copy-number alteration calling

Somatic copy-number variants (CNVs) were called using the Allele-Specific Copy number Analysis of Tumours (ASCAT) algorithm^58^, https://github.com/Crick-CancerGenomics/ascat) in the ascatNGS package^59^. Bulk blood samples or phylogenetically unrelated normal samples were used as matched normals. ASCAT was initially run with default parameters. To reduce the number of false-positive calls that arise when analysing normal tissue samples using ASCAT, a bespoke algorithm ascat-PCA was applied. ascat-PCA extracts a noise profile by aggregating the LogR ratio from across a panel of normal unrelated samples and subtracts this signature from that observed in the sample being analysed using principal component analysis (INSERT GITHUB).

### Structural variant calling

Whole-genome sequences were analysed for somatic structural variants (SVs) using the Genomic Rearrangement Identification Software Suite (GRIDSS). In preparation for this analysis, genomes were remapped to Human Genome Version 38 and GRIDSS was run using the same matched normal as used for CNV analysis. Coordinates for SV calls were subsequently converted back to GRCh37. SV calls in L1 transposon donor regions and fragile sites were excluded from the final SV analysis.

### Mutational signature analysis

The R package HDP (https://github.com/nicolaroberts/hdp), based on the hierarchical Dirichlet process^60^, was used to extract mutational signatures. Analysis of mutational signatures using this package has been applied to normal tissues previously^1,4^. In brief, this nonparametric Bayesian method models categorical count data using the hierarchical Dirichlet process. A hierarchical structure is established using patients as the first tier (parent nodes) and individual samples as the second tier (dependent nodes). Uniform Dirichlet priors were applied across all samples. The algorithm creates a mutation catalogue for each sample and infers the distribution of signatures in any one sample using a Gibbs sampler. We performed mutational signatures analysis per-branch, counting each branch of the phylogenetic tree as a distinct sample to avoid double counting of mutations. Since the MCMC process scales linearly with the number of counts, we randomly subsampled each branch to a maximum of 2500 total substitutions. Branches with fewer than 100 mutations were excluded from the mutational signature extraction. No reference signatures were included as priors.

To assess the contribution of each mutational process, mutational signatures were refitted to all mutation counts of branches of phylogenies using the R package sigfit (https://github.com/kgori/sigfit)^61^. To avoid overfitting, a limited subset of reference mutational signatures were included per patient corresponding to the HDP signatures that have been identified in that individual.

Ageing signatures SBS1 and SBS5 are present in all normal intestinal crypts^3^. Lower than expected burdens of SBS1 and SBS5 were observed in most individuals in this study due to; 1. the inherent challenges of accurately estimating mutation burden in hypermutated samples and 2. the appreciable contamination of reference signatures with SBS1 and SBS5. To partially address this, we used the extracted HDP component corresponding to SBS36 in the refitting stage which has lower SBS1 and SBS5 contamination than the COSMIC reference SBS36 signature. Nevertheless, in individual PD44890 where SBS18 and SBS36 exposures are many tens of times greater than the normal mutation rate, the estimates of SBS1 and SBS5 are substantially lower than would be expected.

### Cancer driver mutations

Cancer driver mutations were identified using two methods aiming to identify genes and mutations in this cohort that are subject to positive selection. Firstly, to identify mutations in cancer genes under positive selection in an unbiased manner, we ran a modified dNdS method^62^. To avoid double-counting of mutations, only unique mutations (SBS and ID) which were mapped to branches of the phylogenetic trees were analysed. dNdScv was run using the following parameters; max_coding_muts_per_sample=5000 and max_muts_per_gene_per_sample=20. Genes with a qval of <0.05 were considered to be under positive selection.

A second phase of cancer gene mutation analysis was undertaken; identifying mutations in this cohort which are codified in cancer mutation databases and exhibit characteristic traits of cancer driver mutations; an approach previously employed in the study of normal tissues^1,2^. In this phase of the analysis we sought to identify the spectrum and frequency of cancer driver mutations in this cohort. Somatic mutations (SBS and ID) were collated per-sample from all tissues. Analysis was restricted to protein coding regions and mutations were filtered using lists of known cancer genes; mutations in samples from intestinal epithelium were filtered using a list of 90 genes associated with colorectal cancer and includes variants that are commonly identified in small bowel adenocarcinoma^3^; samples from all other tissues including blood were filtered using a pan-cancer list of 369 driver genes^62^. Genes were then characterised according to their predominant molecular behaviour; dominant, recessive or intermediate (those demonstrating aspects of both types of behaviour) using the COSMIC Cancer Gene Census^63^. All candidate mutations were annotated using the cBioportal MutationMapper database (https://www.cbioportal.org/mutation_mapper). Mutations meeting the following criteria were considered to be driver mutations; truncating mutations (those that cause a shortened RNA transcript, nonsense, essential splice-site, splice region and frameshift ID) in recessively acting genes, known activating hotspot mutations in dominant (and recessive) genes and lastly mutations that were in neither of the above categories but characterised by the MutationMapper database as being ‘likely oncogenic’ were also included in the final driver mutation catalogue. We also sought to compare the frequency of driver mutations in histologically normal crypts with MUTYH mutations to those from individuals who do not carry DNA polymerase mutations. Somatic mutations from 445 normal intestinal crypts^3^ were annotated and filtered using the above criteria. Comparison was made with normal intestinal crypts from this cohort of individuals with MUTYH germline mutations.

### Modified duplex sequencing (NanoSeq)

DNA from bulk blood samples from individuals with germline *MUTYH* mutations was extracted as outlined above. Samples from normal healthy control was obtained and processed using the following method. Whole blood was diluted with PBS and mononuclear cells (MNC) were isolated using lymphoprepTM (STEMCELL Technologies) density gradient centrifugation. The red blood cell and granulocyte fraction of the blood was then removed. The MNC fraction was depleted of red blood cells by lysis steps involving 3 incubations at room temperature for 20 mins/10 mins/10 mins respectively with RBC lysis buffer (BioLegend). Tissue lymphocytes were isolated from Peyer’s patches in intestinal mucosa using laser capture microdissection and subjected to protein lysis as outlined above. Cell lysates were processed and whole genome sequenced using the NanoSeq protocol.

Our modified duplex sequencing method, called NanoSeq, relies on blunt-end restriction enzymes to fragment the genome in order to avoid errors associated to the filling of 5’ overhangs and the extension of internal nicks during end repair after sonication. Our modified method has error rates < 5e-9^50^.

Given the uneven frequencies of trinucleotides in the digested genome, the strong filtering of common SNPs sites (typically occurring at CpG), and the strong dependence of mutation rates on trinucleotide contexts, our estimates of mutation burdens are normalized and projected onto genomic trinucleotide frequencies.

Let *t* denote the count of a given trinucleotide of type *i* = 1…32. The frequency of each trinucleotide is calculated separately for the genome 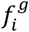 and for the NanoSeq experiment 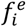 where:

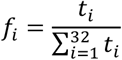

The ratio of genomic to experimental frequencies for a given trinucleotide is:

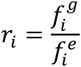

There are *j* = 1…6 classes of substitution where the mutated base is a pyrimidine. Let *s_ij_* denote the count of substitution *j* in trinucleotide context *i*, giving a total of 96 substitution classes. Each substitution count is corrected as follows:

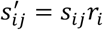

The corrected substitution counts provide a substitution profile projected onto the human genome, and are also used to calculate the corrected mutation burden:

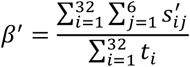

Software used in this study is publicly available at the following locations. Mutation calling algorithms are available at https://github.com/cancerit. Code for filtering mutation calls is available at https://github.com/TimCoorens/Unmatched_NormSeq. Software for mutational signature analysis is available at https://github.com/nicolaroberts/hdp, https://github.com/kgori/sigfit and https://github.com/AlexandrovLab. Software for analysis of duplex / NanoSeq data is provided at https://github.com/cancerit/NanoSeq. Parameters used for these various pieces of software have been included in the manuscript methods section and supplementary information.

## Data Availability

DNA sequencing data are deposited in the European Genome-Phenome Archive (EGA) with accession code: EGAD00001007958 and EGAD00001007997.

The cBioPortal MutationMapper database was accessed at: https://www.cbioportal.org/mutation_mapper?standaloneMutationMapperGeneTab=ATM

## Code Availability

Code/software required to reproduce the analyses in this paper are available online and are listed in Methods.

## ACKNOWLEDGEMENTS

We thank the staff of Wellcome Sanger Institute Sample Logistics, Genotyping, Pulldown, Sequencing and Informatics facilities for their contribution including Laura O’Neill, James Hewinson, Yvette Hooks, Stephen Gamble, Calli Latimer and Kirsty Roberts for their support with sample management and laboratory work. In addition; Moritz Gerstung and Harald Vöhringer for help with analysis, advice and discussions. James Chan (Cambridge University Hospitals) and Geraint Williams (Cardiff University) for assistance with histopathological review. We thank the clinical and research administration teams at recruitment sites for their assistance with sample collection and patients and their families for their time as study participants.

## AUTHOR CONTRIBUTIONS

P.S.R., M.R.S., J.R.S. and L.T. conceived the study design. J.R.S., L.T., F.L., L.B., N.L., H.D.W., R.B-S., S.R, R.t.H., N.C., S.J.A.B. and K.S.P. recruited individuals, collected samples and curated sample and clinical data. P.S.R., B.C.H.L. and S.V.L. undertook laboratory work. F.A., I.M., S.V.L., L.M.R.H., T.H.H.C. and M.A.S. developed bespoke DNA library preparation, sequencing and bioinformatic methods. F.A., I.M., L.M.R.H., H.L.S. and S.O. contributed and analysed control data. P.S.R., F.A., I.M., S.O. and H.J. performed data analysis. M.R.S., P.J.C. and I.M. oversaw statistical analysis. M.R.S. and J.R.S. oversaw the study. All authors were involved in the preparation and review of the manuscript.

